# Lateralized decrease of parvalbumin^+^ cells in the somatosensory cortex of ASD models is correlated with unilateral tactile hypersensitivity

**DOI:** 10.1101/2020.09.08.288654

**Authors:** Tara Deemyad, Stephanie Puig, Andrew Papale, Hang Qi, Gregory M LaRocca, Deepthi Aravind, Emma LaNoce, Nathaniel N Urban

**Affiliations:** Department of Neurobiology, University of Pittsburgh, Pittsburgh, PA; Department of Psychiatry, Translational Neuroscience Program, University of Pittsburgh, Pittsburgh, PA; Center for Neuroscience at the University of Pittsburgh, University of Pittsburgh, Pittsburgh, PA

**Author notes:** Corresponding author: Tara Deemyad.

## Abstract

Inhibitory control of excitatory networks contributes to cortical functions. Increasing evidence indicates that parvalbumin expressing (PV^+^) basket cells (BC) are a major player in maintaining the balance between excitation (E) and inhibition (I) in the cortex. Disruption of E/I balance in cortical networks is believed to be a hallmark of autism spectrum disorders (ASD) and may contribute to sensory alterations seen in ASD. Here, we report a lateralized decrease in the number of PV^+^ BCs in L2/3 of the somatosensory cortex in the dominant hemisphere of adult Shank3^-/-^ and Cntnap2^-/-^ mouse models of ASD. The dominant hemisphere was identified during a reaching task to establish each animal’s dominant forepaw. Double labeling with anti-PV antibody and a biotinylated lectin (i.e., VVA) showed that the number of BCs was not different but rather, some BCs did not express detectable levels of PV (PV^-^), resulting in an elevated number of PV^-^ VVA^+^ basket cells. This lateralized reduction was not observed in the number of interneurons from the other two major groups that express somatostatin or the serotonergic receptor 5HT3a. Finally, we showed that dominant hind paws had higher mechanical sensitivity (i.e., lower mechanical thresholds measured with von Frey test) but no difference in thermal sensitivity (measured with Hargreaves test) when compared to the other hind paw. This mechanical hypersensitivity in the dominant paw correlated with the decrease in the number of PV^+^ interneurons and reduced PV expression in the corresponding cortex. Together, these results suggest that the sensory hypersensitivity in ASD could be due to decreased inhibitory inputs to the dominant somatosensory cortex.

## Introduction

Autism spectrum disorder (ASD) is a neurodevelopmental disorder that is described by a broad range of cognitive and behavioral abnormalities. A wide range of environmental, genetic, and biological factors have been implicated in the pathophysiology of ASD (Mullins C et al. 2016). Reduction in signal to noise ratio due to disruption in balance between excitatory and inhibitory inputs or E/I imbalance in neural circuits is an influential hypotheses that provides a common neural basis of ASD (Rubenstein JL and MM Merzenich 2003). This hypothesis has been supported by clinical and biological analyses over the past two decades (Oblak AL et al. 2010; Chattopadhyaya B and GD Cristo 2012; Lee E et al. 2017; Masuda F et al. 2019). In particular, growing evidence suggests that E/I imbalance in ASD subjects arises from dysregulation of cortical GABAergic interneurons (Pizzarelli R and E Cherubini 2011; Martin BS et al. 2014; Dickinson A et al. 2016).

Cortical interneurons can be divided into three non-overlapping groups based on expressing parvalbumin (PV), somatostatin (SST) or ionotropic serotonin receptor 5HT3a (5HT3aR) (Rudy B et al. 2011). Parvalbumin positive interneurons, particularly basket cells (BC), have been investigated in more detail and previous studies suggest that they are critical for maintaining E/I balance (Ferguson BR and WJ Gao 2018). Different studies have shown a decrease in the number of PV-immunoreactive (PV^+^) BCs or a reduction in PV expression in BCs in the cortex of both individuals with ASD (Ariza J et al. 2018; Hashemi E et al. 2018; Selten M et al. 2018) and mouse models of ASD (Filice F et al. 2016; Lauber E et al. 2018; Vogt D et al. 2018). Furthermore, it has been shown that an increase in the activity of excitatory neurons could also result in E/I imbalance and ASD-like symptoms, such as produced in the valproic acid (VPA) model of ASD (Kim KC et al. 2013). Finally, restoring the E/I balance in ASD mouse models through enhancing GABAergic activity (Han S et al. 2012; Han S et al. 2014) or inhibiting glutamatergic activity (Kim JW et al. 2017) can rescue ASD-like behaviors.

Interneurons play a crucial role in modulation of brain activity and in functions such as cognitive flexibility, sensory interpretation (e.g., touch, vision and hearing), attention, and social interaction (Deemyad T et al. 2018; Mederos S and G Perea 2019). Emerging data suggest that BCs play an important role in structuring the temporal properties (i.e., rhythmic pacing) and coordination of neuronal responses (Royer S et al. 2012; Tzilivaki A et al. 2019) in oscillating networks. Such synchronization of activity between different cortical areas as well as between the two hemispheres is an important feature of normal brain development and function (Ermentrout GB et al. 2008; Uhlhaas PJ et al. 2008) and could be a common source for the wide range of symptoms observed in ASD.

Imbalance in BCs between the two hemispheres and the resultant asymmetric E/I imbalance would be expected to have a powerful impact on sensory processing and cognition. Studies on ASD subjects have suggested hemispheric abnormalities in microstructure and function (Dawson G et al. 1983; Perkins TJ et al. 2014; Peterson D et al. 2015). For instance, clinical studies suggest that ASD subjects may show lateralized deficits in steady-state gamma responses (Wilson TW et al. 2007). Furthermore, two mouse models of ASD (VPA and neuroligin-3 mutation) show interhemispheric asymmetry in the number of BCs (Gogolla N et al. 2009).

Here we describe a series of experiments in which we investigated the distribution of the three main classes of GABAergic interneurons in dominant and non-dominant somatosensory cortices (identified after performing a behavioral reaching task) of adult mice in two models of ASD in which autism candidate genes contactin associated protein-like 2 (Cntnap2^-/-^) or SH3 and multiple ankyrin repeat domains 3 (Shank3^-/-^) were deleted. We found a robust decrease in both the number of BCs and their PV expression with no change in the other two classes of GABAergic interneurons (i.e., 5HT3a and SST). Notably, this change was limited to the dominant hemisphere. This lateralization in BCs was correlated with a hypersensitivity to mechanical stimulation of the dominant paw. These findings suggest that a decrease in inhibitory cortical inputs can be a possible neural mechanism for the hypersensitivity observed in subjects with ASD.

## Materials and Methods

### Animals

All procedures were performed in accordance with protocols approved by the University of Pittsburgh Institutional Animal Care and Use Committee (IACUC). Data were obtained from a total of 19 mice (male and female). C57BL/6 mice were obtained from Charles River Laboratories; all others (Cntnap2^+/-^, Cntnap2^-/-^, Shank3^+/-^ and Shank3^-/-^) were obtained from Jackson Laboratories. Mice from the same litter but different genotypes, i.e., littermate controls, are included.

### Immunohistochemistry

Mice were given an overdose of ketamine-xylazine. Once non-responsive, they were perfused transcardially by gravity perfusion with a 20-gauge needle for one minute with cold 0.1 M phosphate-buffer (PB) containing 1% NaCl, followed by cold 4% paraformaldehyde in PB (PFA). (Approximate volume of each was 35 ml.) Brains were removed and fixed in PFA overnight at 4 °C, and then were transferred to 30% sucrose in PB for cryoprotection. Coronal slices (25 µm) were cut using a sliding microtome (Leica SM2010R) and maintained at 4C in PB containing 0.05% Sodium Azide and 0.005% Tween 20 before immunochemistry and/or mounting. The slices covered the entire somatosensory cortex according to Allen Atlas (Fig. 1A).

**Figure 1.**
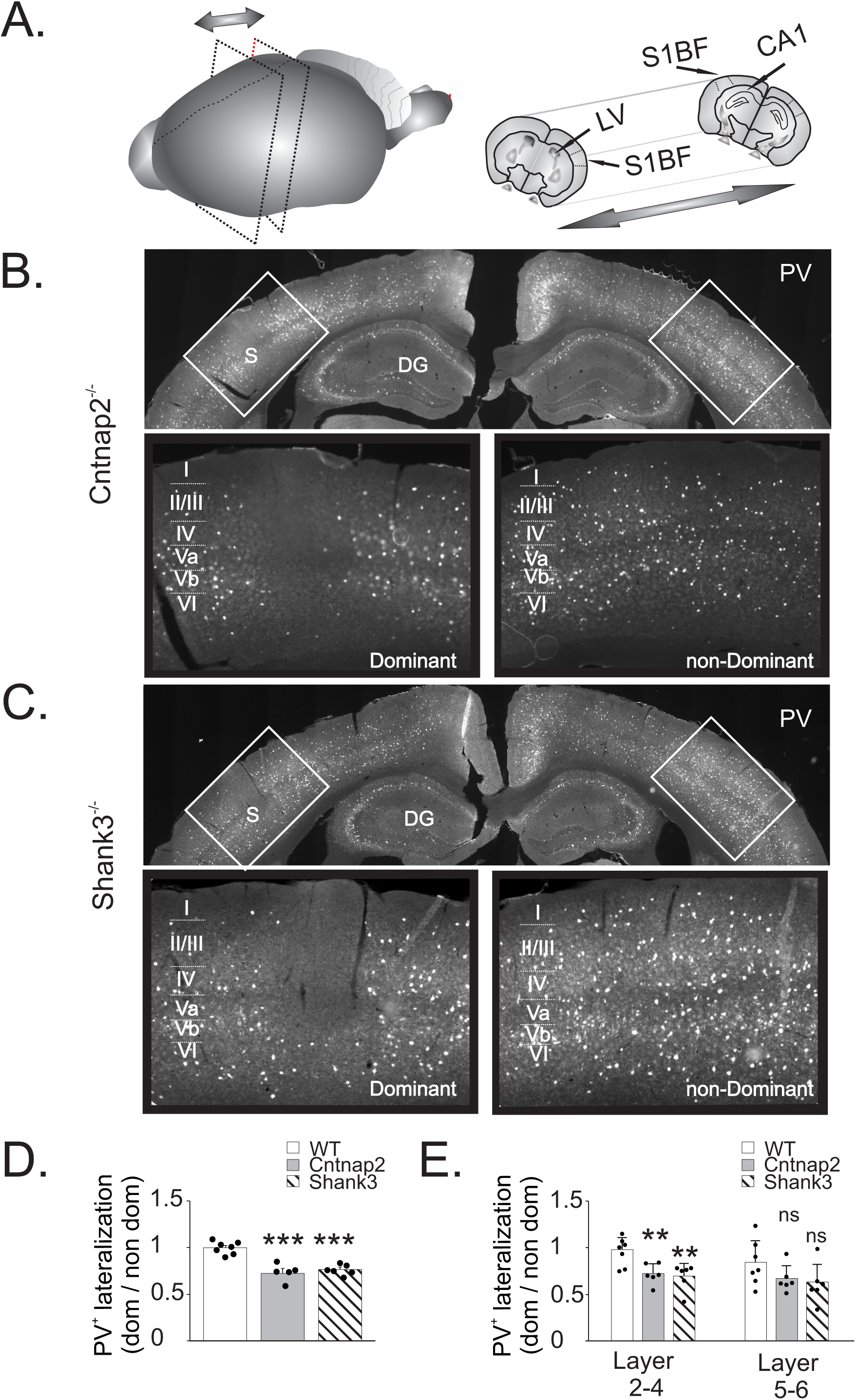
PV interneurons are reduced in dominant hemisphere of Shank3^-/-^ and Cntnap2^-/-^ mice. (A) Schematic of the location of the cortical patches used in PV staining. (B) Representative PV immunofluorescence images of Cntnap2^-/-^ mouse line (top) and ROIs (bottom) of dominant (left) and non-dominant (right) hemispheres. (C) Representative PV immunofluorescence images of Shank3^-/-^ mouse line (top) and ROIs (bottom) of dominant (left) and non-dominant (right) hemispheres. (D) Significant differences are observed in PV^+^ cell count lateralization ratio in Shank3^-/-^ and Cntnap2^-/-^ mice compared to WT mice. (E) Significant differences are observed in PV^+^ cell count lateralization ratio in layers II/III of Shank3^-/-^ and Cntnap2^-/-^ compared to WT. *Significant vs. WT mice. Asterisks represent *P ≤ 0.05, **P ≤ 0.01, ***P ≤ 0.001, respectively. Data are from 5 (Cntnap2^-/-^) and 6 (Shank3^-/-^)independent experiments. Error bars represent SEM.

Free-floating sections were permeabilized and blocked by incubation in 0.1% Triton X-100 and 2% normal donkey serum (NDS; Jackson ImmunoResearch #017-000-121) in PB for 1 hour. Sections were washed 3 times with PB for 5 minutes using oscillation at RT. This washing procedure was performed following each antibody or labeling step. Sections were then mounted on collagen coated slides in gelvatol and imaged with Nikon Eclipse 90i and E600 microscopes.

Antibodies were added at the indicated concentrations in 2% NDS and 0.05% Tween 20 in PB. Sections were labeled with either anti-parvalbumin or anti-5HT3A receptor. Sections labeled with mouse anti-parvalbumin (PV; Sigma #P3088) were incubated at 1:1000 overnight at 4C. This was followed by Alexa Fluor 488 donkey anti-mouse antibody (ThermoFisher Scientific #A-21202) at 1:600 and Hoechst 33342 (ThermoFisher Scientific #H3570) at 1:40,000 for 1 hour with oscillation at RT. In some cases, sections labeled with PV were then incubated in biotinylated Vicia Villosa Lectin (VVA; Vector Laboratories #B-1235) at 1:400 in PB for 2 hours with oscillation at RT. This was followed by Alexa Fluor 594 Streptavidin (ThermoFisher Scientific #S32356) at 1:4000 in PB for 1 hour with oscillation at RT.

For anti-5HT3A labeling, sections were incubated with 1:400 rabbit anti-5HT3A antibody (HTR3A; Alomone Labs #ASR-031) at 4°C for two days. This was followed by Alexa Fluor 594 donkey anti-rabbit antibody (ThermoFisher Scientific #A-21207) at 1:600 and Hoechst 33342 at 1:40,000 for 1 hour, with oscillation at RT.

### Automated image analysis

All data analysis was performed in MATLAB (The MathWorks, Natick, MA) and Fiji (Schindelin J et al. 2012). Specifically, a previously described MATLAB based command line software toolbox, cell segmentation (CellSegm)(Hodneland E et al. 2013) was used for counting surface stained cells (VVA and SST staining; Hodneland E et al. 2013). A custom-written MATLAB code using simple thresholding was used for counting cytoplasmic staining (PV staining). Lateralization ratio for each measurement was defined by the measured value of dominant / non-dominant hemisphere in each animal. Cells were counted in matched areas of fixed size in each hemisphere of somatosensory cortex.

### Behavioral procedures

Animals were randomly allocated to the different groups. To decrease bias toward the outcome of the experiment, researchers who were running and analyzing these experiments were blind to this allocation.

#### Food retrieval task of forelimb dominance (reaching task)

Mice of at least 6 weeks old were trained for three days before performing a reaching task (Chen CC et al. 2014). Briefly, after measuring baseline bodyweight for two days prior to behavioral test, mice were food-restricted to initiate bodyweight loss. The amount of food was adjusted based on baseline bodyweight, to 10% of the original baseline weight. Next, mice were habituated as a group for two days. For this step, two mice were placed in a training chamber. About 20 seeds/mouse were placed inside the chamber for their intake. Mice stayed in the chamber for 20 min and then put back into their home cage. On the third day, mice were habituated separately by placing mice into the training chamber individually with access to 20 seeds for 20 min. On the testing day, the food tray was filled with seeds and advanced against the front wall of the training chamber to allow the seeds to be available to the mouse (movie. S1). Critically, 20 or more reaches with one forelimb comprising 70% of the total reaches during each 5-minute session, was required for a successful trial (Chen, Gilmore et al. 2014). If the mouse used its tongue to get the seeds into the chamber, the tray was moved back from the slit slightly. Each trial was videotaped and the number of reaches of each paw counted off-line by an investigator who was blind to the experimental conditions.

#### Hargreaves test

Mice were individually placed in Plexiglas chambers on the tempered glass of a Hargreaves apparatus warmed at 30° C (IITC Life Science Inc., Woodland Hills CA), and allowed to habituate. A focused light beam was applied to the plantar surface of left and right hind paws alternatively (Fig. 5A schematic). The duration of the stimulus that resulted in paw withdrawal was considered as the paw withdrawal latency (PWL). Three different light intensities inducing low, medium or high thermal stimulations (set at 10%, 15% and 40% respectively of the maximum possible intensity of the Hargreaves apparatus) were used. A cutoff of 20 seconds was used to avoid tissue damage. The mean of four measures of PWLs applied at intervals of 5 minutes, was determined for each light intensity (Saloman JL et al. 2016).

#### von Frey mechanical test

Mice from different strains were individually placed in Plexiglas chambers on a wire mesh grid and were allowed to habituate (Fig. 6A schematic). The plantar surface of the hind paws was stimulated using von Frey filaments. We used 3 different forces: 2.83 g, 3.22 g, and 4.08 g defined as weak, medium and strong force mechanical stimuli, respectively. Each filament was directed to the center of the plantar surface of the targeted paw and gradually pressed upward until the filament bent and was then held for 2 seconds. Left and right hind paws were stimulated 10 times with each filament, with 5 min intervals between stimuli. A response was considered positive if the animal withdrew, bit, shook or licked the stimulated paw during the 2 s of stimulation. Otherwise, the response was taken as negative (Cui L et al. 2016). The percentage of positive responses was calculated and reported as previously described in (Saloman JL et al. 2016).

### Statistics

All measurements are presented as mean ± SEM unless otherwise indicated. For comparison between two conditions in the same animal paired Student’s t-test or repeated measures two-way ANOVA with Tukey’s or Sidak’s post hoc test was used. Statistics were analyzed using Prism 7.

## Results

### Asymmetric reduction in PV^+^ cells in the somatosensory cortex of Cntnap2^-/-^ and Shank3^-/-^ mice

Previous studies have shown decreased numbers of GABAergic neurons and in particular PV^+^ subgroup by ∼20–25% in several ASD mouse models and subjects (Gant JC et al. 2009; Gogolla N *et al*. 2009; Tripathi PP et al. 2009; Wohr M et al. 2015; Hashemi E *et al*. 2018). Furthermore, an asymmetric reduction of cortical PV^+^ cells in VPA and NL-3^-/-^ mouse models of ASD have been reported (Gogolla N *et al*. 2009). If similar changes are observed in different animal models of ASD then it suggests a common mechanism for shared overarching symptoms. As an initial step, we explored whether such asymmetry in the number of PV^+^ interneurons is also present in Shank3^-/-^ (n =6) and Cntnap2^-/-^ (n = 5) models of ASD. We observed an asymmetry in the number of PV^+^ cells between the two hemispheres that was clear even at low magnification in both of these mouse models (Fig. 1B). Overall, the average number of PV^+^ cells in both hemispheres were 91±10, 90±4, and 131±7 cells/mm^2^ in the somatosensory cortex of Cntnap2, Shank3 and WT mice, respectively This reduction was similar to previous studies (Gogolla N et al. 2014; Filice F *et al*. 2016; Vogt D *et al*. 2018). However, the ratio of PV^+^ cell counts in the L/R hemispheres was not significantly different between groups. In fact, the reduction in PV^+^ cells in each animal was restricted to one hemisphere, but which hemisphere (L vs R) was not consistent across animals. Such reduction in PV^+^ cells was not also uniform among cortical regions and slices. This reduction seemed to be most pronounced in the primary and supplemental somatosensory cortical areas, followed by the dorsal and primary auditory cortices (Figs. 1B and 1C). Interestingly, the somatosensory system is highly affected in ASD and alterations in somatosensation are correlated to the severity of ASD (Tomchek SD and W Dunn 2007; Wiggins LD et al. 2009; Mammen MA et al. 2015; Orefice LL et al. 2019). Therefore, we mainly focused on the somatosensory cortex and the side of this lateralized decrease in PV^+^ neurons.

Different mice showed fewer PV^+^ cells in either the right (36% of mice) or left hemisphere (63% of mice).We investigated whether the affected side had any relation to the dominant hemisphere based on the handedness of the animal. We used a procedure similar to that described by a previous study for determining paw dominance in mice (Waters NS and VH Denenberg 1994). Nineteen adult mice were trained for a reaching task in which mice were placed in a plexiglass chamber with access to small seeds through a slit small enough to fit a single forepaw (Supplementary Movie1). Both Shank3^-/-^ and Cntnap2^-/-^ mice became familiar with the task in less than a week, with 50% of them showing left paw preference. One of the animals was ambidextrous and was excluded from the study. Cortical slices from these animals were then immunostained for PV and the number of cells in the somatosensory cortex was counted. To compare the two hemispheres, we used the ratio of the number of PV^+^ cells in the dominant hemisphere (i.e., opposite to the dominant paw) over the other hemisphere (Fig. 1D). While this ratio was close to 1 in WT animals (0.99 ± 0.2, n=7), it showed a reduction in both Shank3^-/-^ (0.76± 0.2, ANOVA, p= 0.001, n=6) and Cntnap2^-/-^ mice (0.73±0.3, ANOVA, p= 0.001, n=5), suggesting a reduction of PV^+^ cells in the dominant somatosensory cortex.

We next explored whether this lateralized reduction in PV cell numbers was limited to specific cortical layers. Normally, PV^+^ cells are plentiful throughout layers II-VI and show morphological and functional differences across layers (Tremblay R et al. 2016). In particular axons of PV cells in layers 2/3 and 5/6 extend through several columns, in addition to their local and translaminar innervation (Atallah BV et al. 2012; Tremblay R *et al*. 2016). As reported by other groups (Hafner G et al. 2019), PV^+^ cells were not detected in layer I of the somatosensory cortex in any of the animals. In our samples from ASD and WT groups, PV^+^ cells were present in layers II-IV and V-VI (Figs. 1B and 1C). The most robust lateralization in PV^+^ numbers was observed in layers II-IV (Shank3^-/-^: 0.69 ± 0.05; ANOVA, p=0.01, n =5 and Cntnap2^-/-^: 0.69 ± 0.04, ANOVA, p = 0.01, n =6; vs. WT: 0.96 ± 0.5) and no changes were present in Layer V-VI (Shank3^-/-^: 0.82±0.1; Cntnap2^-/-^: 0.88 ± 0.08, p=0.3; vs. WT:1.1 ± 0.1, p=0.2) (Fig. 1E). Together these results suggest intra- and trans-columnar alteration in inhibitory circuits in the dominant somatosensory cortex of ASD mice.

As a control for the observed asymmetry described above, we also counted PV^+^ cells in the hippocampus. PV^+^ cells were abundant in the stratum pyramidal layer (Fig. 2A and 2B), with a reduction in total numbers in ASD animals, consistent with previous studies (Sik A et al. 1995; Sadakata T et al. 2007; Gant JC *et al*. 2009; Gogolla N *et al*. 2009; Penagarikano O et al. 2011). However, the number of PV^+^ cells were not different between the dominant-side and non-dominant-side CA1 areas (based on handedness) (Shank3^-/-^: 0.79± 0.3, ANOVA, p =0.4, n =6, Cntnap2^-/-^: 0.97± 0.1, ANOVA, p= 0.9, n=5; vs. WT: 1.01 ± 0.4, n=7) or between the left and right CA1 areas (Shank3^-/-^: 1.3 ± 0.4, ANOVA, p = 0.3, n =6 and Cntnap2^-/-^: 1.05 ± 0.2, ANOVA, p= 0.9, n=5; vs. WT: 1.1 ± 0.3, n=7) (Figs. 2C and 2D). Overall, these results suggest that lateralization in inhibitory circuits is specific to certain cortical areas in Shank3^-/-^and Cntnap2^-/-^ mice.

**Figure 2.**
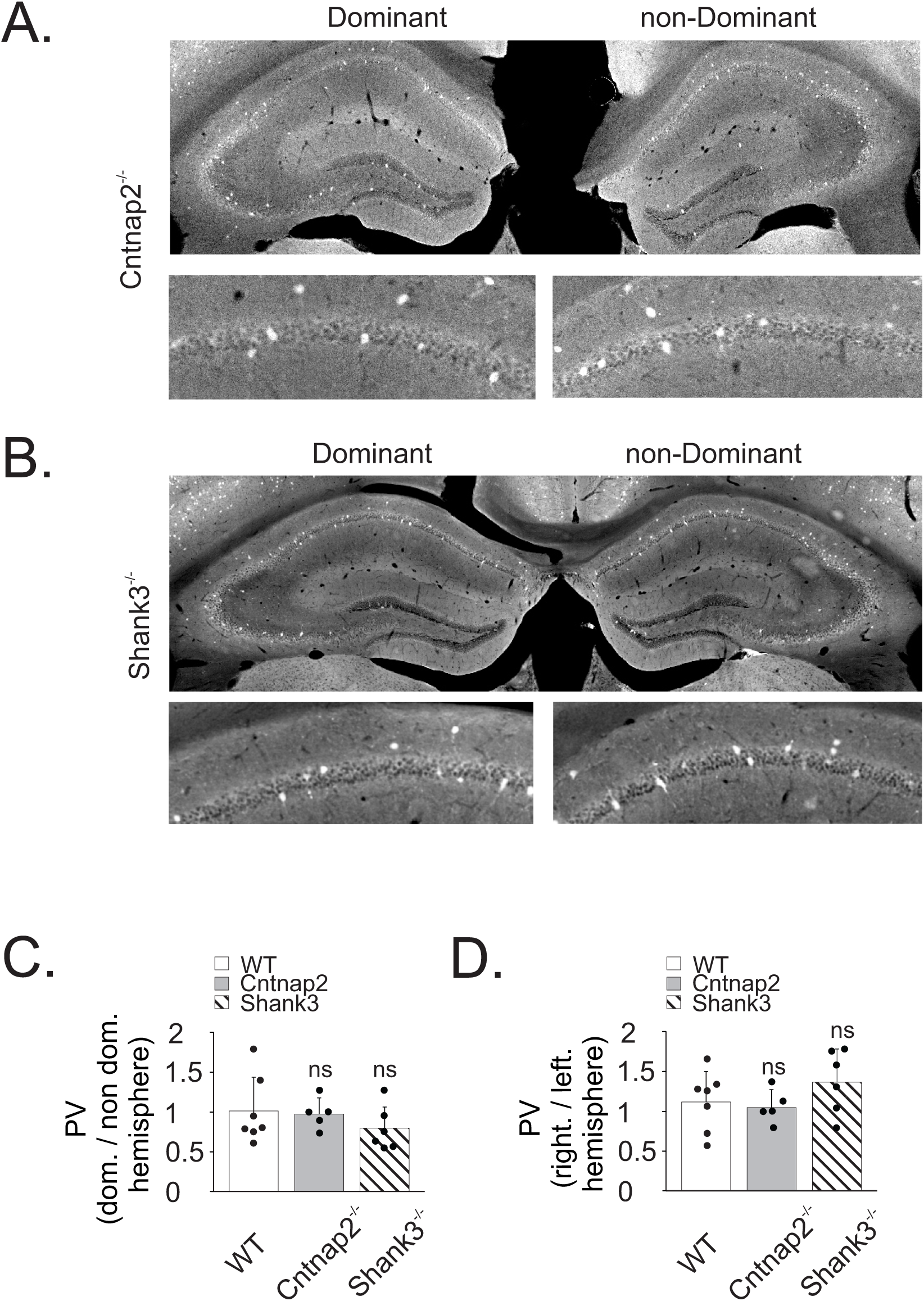
PV interneuron numbers are not changed in hippocampus between two hemispheres of Shank3^-/-^ and Cntnap2^-/-^ mice. (A) Representative PV immunofluorescence images of Cntnap2^-/-^ mouse line (top) and ROIs (bottom) of hippocampus in dominant (left) and non-dominant (right) hemispheres. (B) Representative PV immunofluorescence images of Shank3^-/-^ mouse line (top) and ROIs (bottom) of hippocampus in dominant (left) and non-dominant (right) hemispheres. (C) Lateralization ratio of PV^+^ cell count (dominant/non-dominant) is not significantly different in Shank3^-/-^ and Cntnap2^-/-^ compared to WT. (D) Lateralization ratio of PV^+^ cell count (right/left hemishperes) is not significantly different in Shank3^-/-^ and Cntnap2^-/-^ compared to WT. Error bars represent SEM.

**Figure 3.**
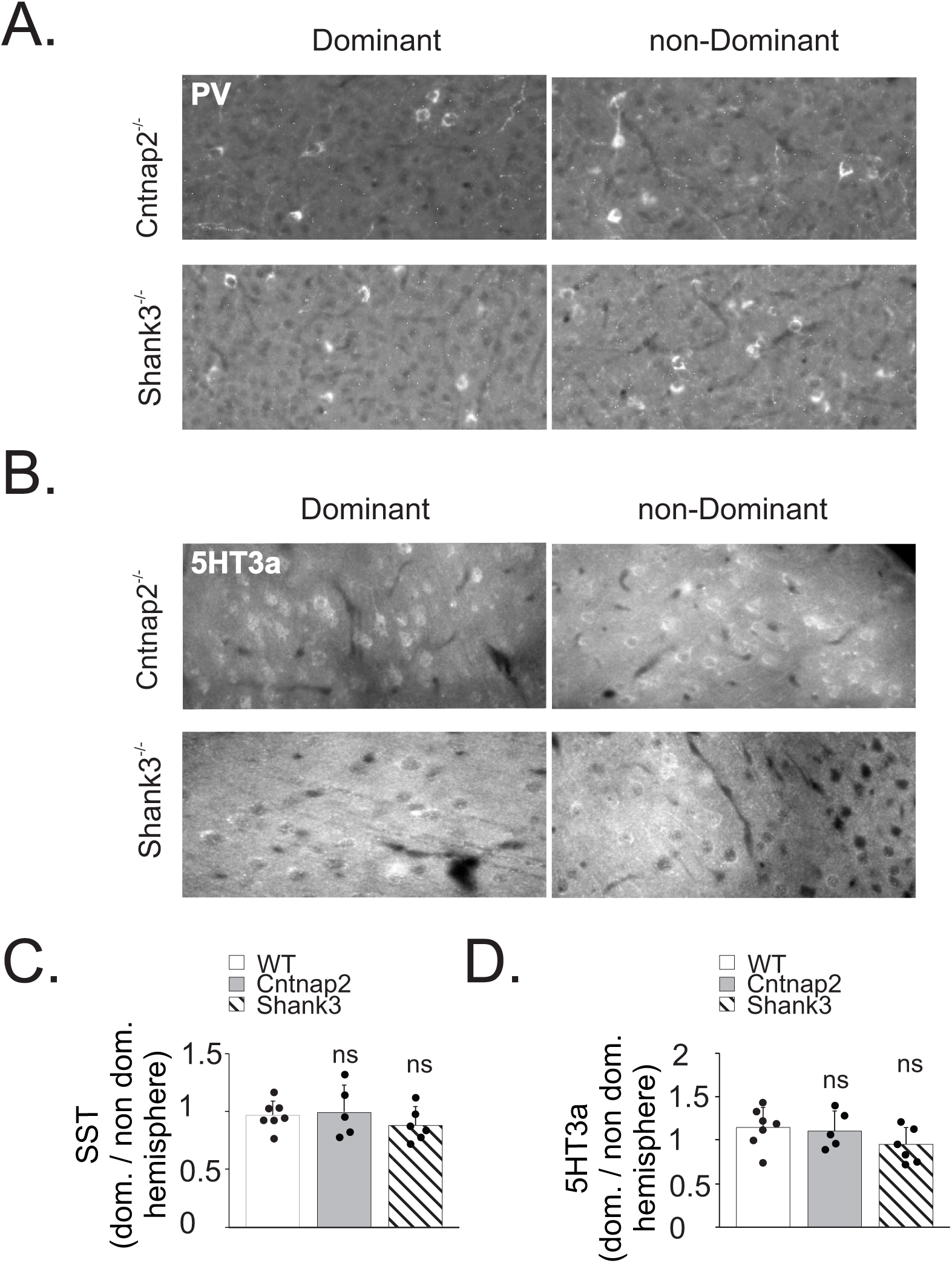
5HT3a and SST interneurons cell counts are not changed between somatosensory cortices of Shank3^-/-^ and Cntnap2^-/-^ mice. (A) Representative SST immunofluorescence images of Cntnap2^-/-^ (top) and Shank3^-/-^ (bottom) mice lines of somatosensory cortex in dominant (left) and non-dominant (right) hemispheres. (B) Representative 5HT3a immunofluorescence images of Cntnap2^-/-^ (top) and Shank3^-/-^ (bottom) mice lines of somatosensory cortex in dominant (left) and non-dominant (right) hemispheres. (C) Lateralization ratio of SST^+^ cell count (dominant/non-dominant) is not significantly different in Shank3^-/-^ and Cntnap2^-/-^ compared to WT. (D) Lateralization ratio of 5HT3a^+^ cell count (dominant/non-dominant) is not significantly different in Shank3^-/-^ and Cntnap2^-/-^ compared to WT. Error bars represent SEM.

### Number of 5HT3a and SST interneurons in the somatosensory cortex of Cntnap2^-/-^ and Shank3^-/-^ mice are not affected

Previous studies have reported decreased numbers of other GABAergic interneurons in ASD mouse models and postmortem subjects (Penagarikano O *et al*. 2011; Rapanelli M et al. 2017). Here we compared the number of different subgroups of cortical interneuron between the two hemispheres in adult WT mice and ASD mouse models. Expression of PV, somatostatin (SST) or the 5HT3a receptor can be used to classify up to 99% of cortical interneurons in WT mice (Rudy B *et al*. 2011). We used antibodies against SST and 5HT3a in adult Shank3^-/-^, Cntnap2^-/-^ and WT mice and compared the inter-hemispheric ratio in the somatosensory cortex. The inter-hemispheric ratio in ASD and WT mice did not differ for SST^+^ cells (Fig. 4E, Shank3^-/-^: 0.89± 0.06, ANOVA, p = 0.6, n =6, Cntnap2^-/-^: 1± 0.08, ANOVA, p= 0.9, n=5; vs. WT: 0.96 ± 0.05, n=7) or 5HT3a^+^ cells (Fig. 4F, Shank3^-/-^: 0.95± 0.7, ANOVA, p = 0.2, n =6, Cntnap2^-/-^: 1.12± 0.7, ANOVA, p= 0.9, n=5; vs. WT: 1.14 ± 0.1, n=7). These results suggest that only inhibition through PV expressing interneurons is altered in these mice suggesting a unique role for PV^+^ interneurons in the pathophysiology of ASD.

**Figure 4.**
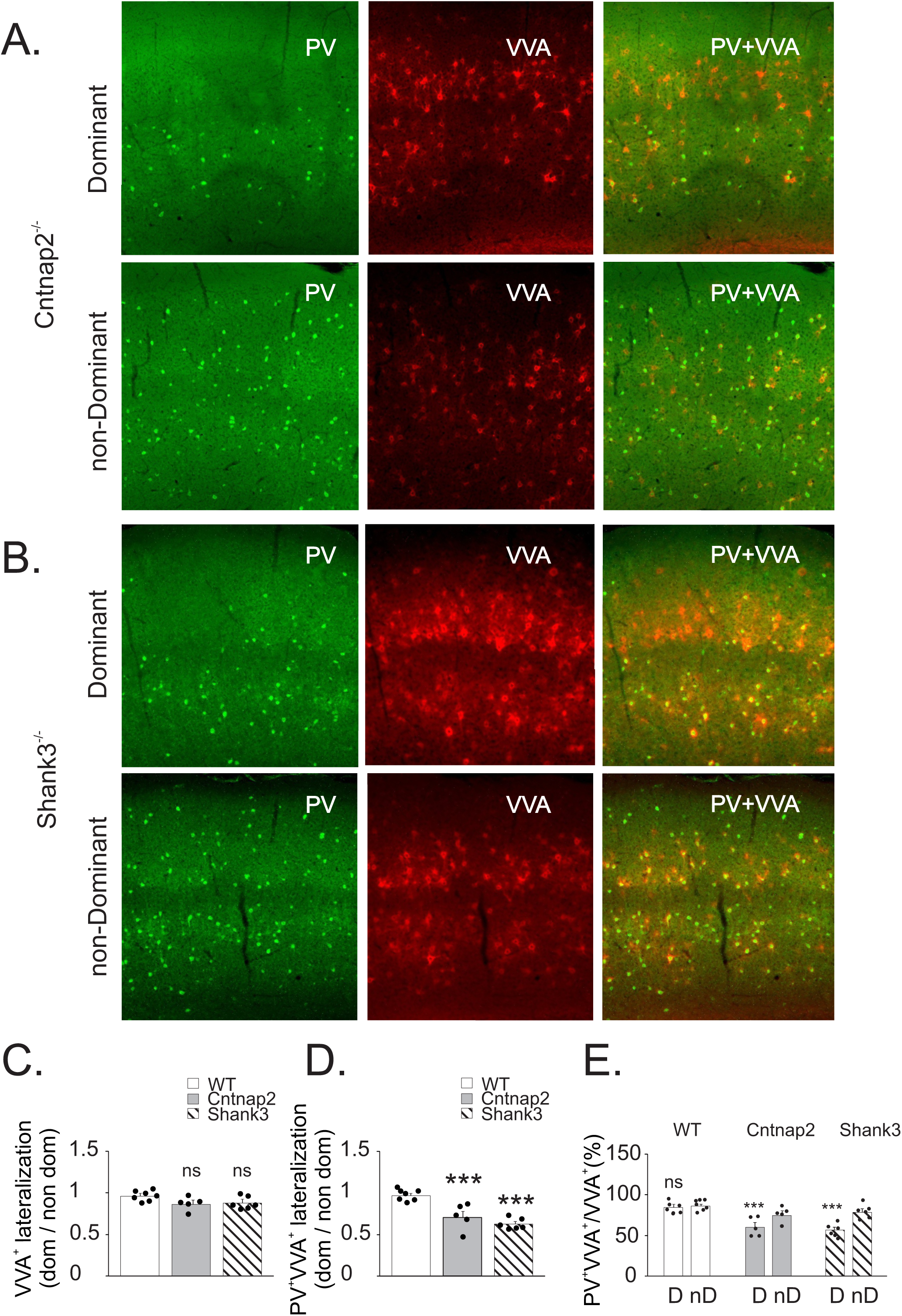
Expression of PV in BC interneurons in the somatosensory cortex is reduced in dominant hemisphere of Shank3^-/-^ and Cntnap2^-/-^ mice. (A) Representative PV immunofluorescence (left), VVA labaled cells (middle) and PV/VVA co-labaled (righ) images of dominant (top) and non-dominant (bottom) hemispheres of Cntnap2^-/-^ mice lines of somatosensory cortex. (B) Representative PV immunofluorescence (left), VVA labaled cells (middle) and PV/VVA co-labaled (righ) images of dominant (top) and non-dominant (bottom) hemispheres of Shank3^-/-^ mice lines of somatosensory cortex. (C) Lateralization ratio of VVA^+^ cell count (dominant/non-dominant) is not significantly different in Shank3^-/-^ and Cntnap2^-/-^ compared to WT. (D) Lateralization ratio of co-labeled VVA/ PV cells (dominant/non-dominant) is significantly reduced in Shank3^-/-^ and Cntnap2^-/-^ compared to WT. (E) Percentage of VVA^+^ cell expressing PV is significantly reduced in dominant hemisphere of Shank3^-/-^ and Cntnap2^-/-^ mice line. Asterisks represent *P ≤ 0.05, **P ≤ 0.01, ***P ≤ 0.001, respectively. Error bars represent SEM.

**Figure 5.**
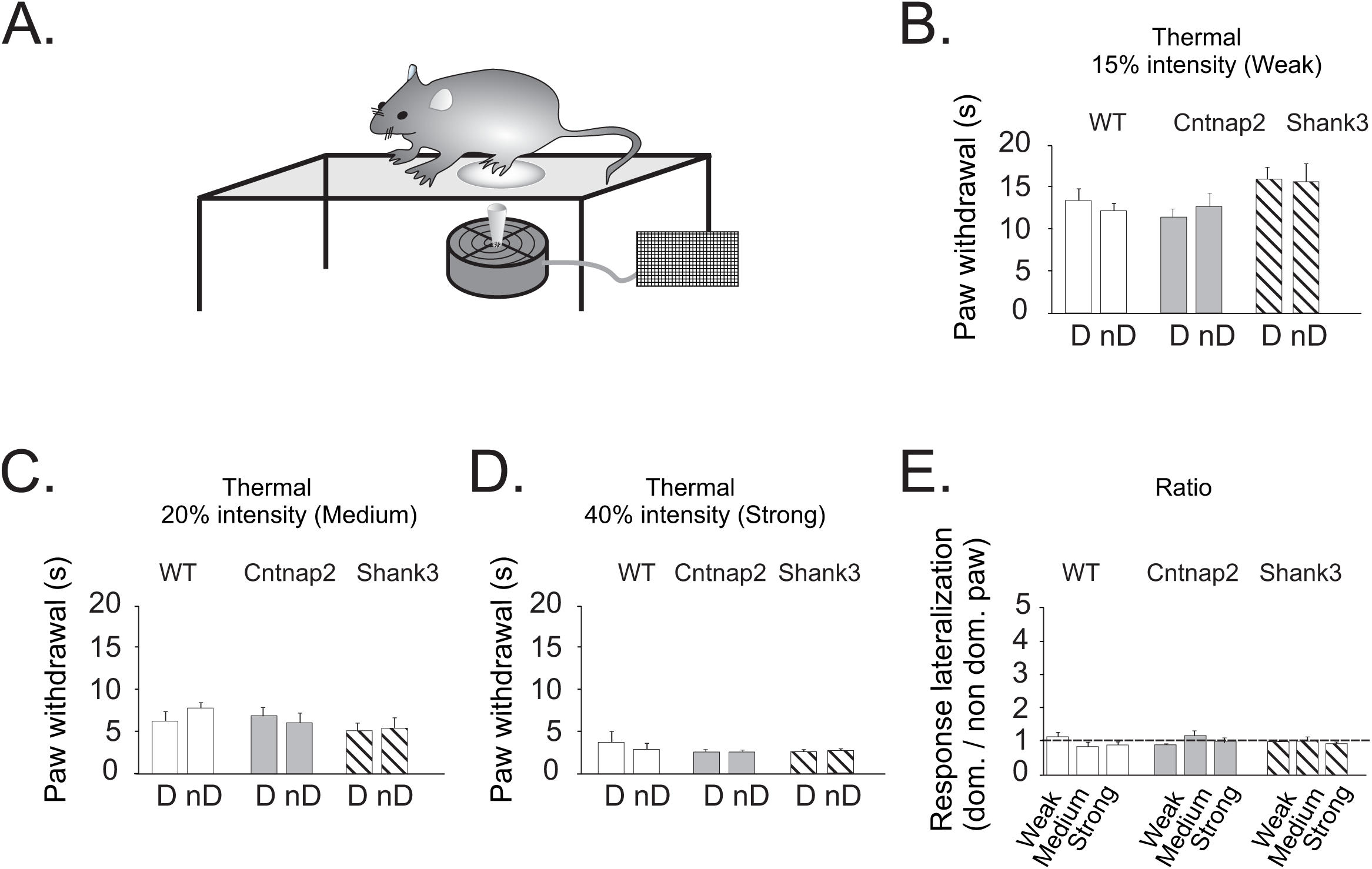
Sensitivity of dominant and non-dominant paws of Shank3^-/-^ and Cntnap2^-/-^ mice to thermal stimuli is not changed. (A) Schematic of the Hargreaves test. Mouse is placed in area with a glass floor. A infrared heat source at three different intensities is focused on the plantar surface of dominant or non-dominant hind paw and the time taken to withdraw from the heat stimulus is recorded for each paw and intensities (B). Withdrawal delay response to weak thermal stimuli is not siginificantly different between dominant and non-dominant of WT, Shank3^-/-^ or Cntnap2^-/-^ mice (C). Withdrawal delay response to medium thermal stimuli is not siginificantly different between dominant and non-dominant of WT, Shank3^-/-^ or Cntnap2^-/-^ mice (D).Withdrawal delay response to strong thermal stimuli is not siginificantly different between dominant and non-dominant of WT, Shank3^-/-^ or Cntnap2^-/-^ mice (E). Response lateralizationis (dominant/non-dominant) is not significantly different between weak, medium and strong thermal stimuli in WT, Shank3^-/-^ or Cntnap2^-/-^ mice. Error bars represent SEM.

**Figure 6.**
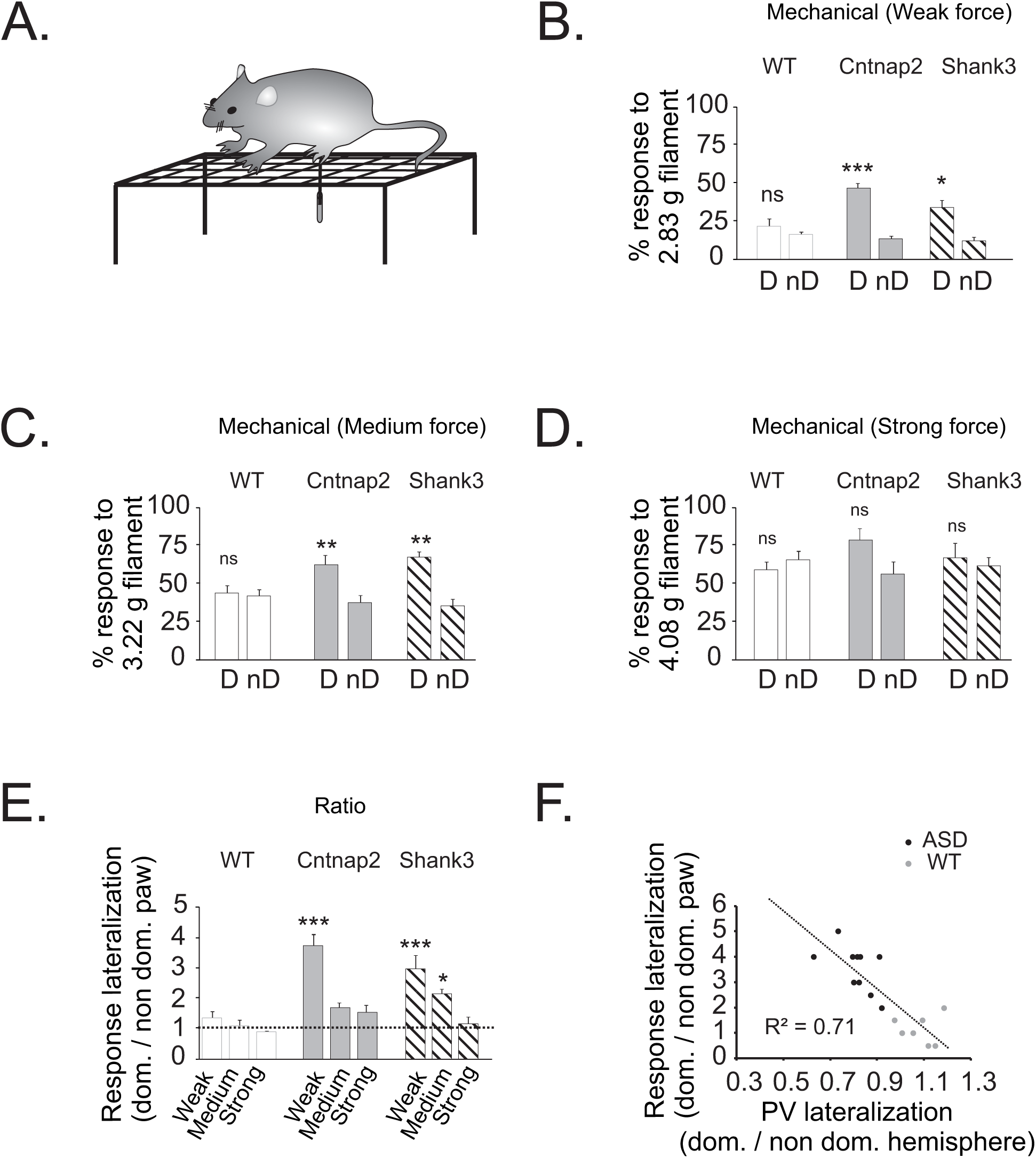
Sensitivity of dominant paws to mechanical stimuli in Shank3^-/-^ and Cntnap2^-/-^ mice is increased. (A) Schematic of manual von Frey test. Mouse is placed on a mesh. Monofilaments of three differing forces are applied perpendicularly to the dominant or non-dominant hind paws. If the rodent withdraws, licks or shakes the paw, it is considered to have had a positive response. (B) The percentage of response to weak mechanical force stimuli by dominant paw in Shank3^-/-^ and Cntnap2^-/-^ mice is significantly higher. (C) The percentage of response to medium mechanical force stimuli by dominant paw in Shank3^-/-^ and Cntnap2^-/-^ mice is significantly higher. (D) The percentage of response to strong mechanical force stimuli by dominant and non-dominant paws of Shank3^-/-^ and Cntnap2^-/-^ mice is not significantly different. (E) Response lateralizationis (dominant/non-dominant) is significantly different in response to weak mechanical force stimuli between in Shank3^-/-^ or Cntnap2^-/-^ and WT mice. (F) Response lateralizationis (dominant/non-dominant) is significantly correlated with PV^+^ cell count lateralization. Asterisks represent *P ≤ 0.05, **P ≤ 0.01, ***P ≤ 0.001, respectively. Error bars represent SEM.

### Decreased expression of PV in BC interneurons in the somatosensory cortex

Recent studies in few ASD mouse models including Shank3^-/-^ and Cntnap2^-/-^ have reported that rather than loss of the interneurons, there is a reduction in expression of PV in basket cells in the cortex and striatum (Filice F *et al*. 2016; Lauber E *et al*. 2018). We studied whether such a decrease in PV expression is more pronounced in the dominant somatosensory cortex. We used Vicia Villosa Agglutinin (VVA) to detect basket cells which are ensheathed by perineuronal nets(Ariza J *et al*. 2018). Double labeling with VVA and anti-PV antibodies was used for identifying basket cells that expressed PV. This also differentiated them from chandelier interneurons that also express PV but are not labeled with VVA. Finally, cells that were labeled with VVA, but not with anti-PV were considered as basket cells that did not express PV (PV^-^ Fig. 4A). We calculated the density (number of cells/mm^2^) of BC interneurons (i.e., density of VVA^+^ cells), and PV (i.e., density of cells double labeled with VVA and PV) for each hemisphere (Fig S1). We did not find a significant reduction in the lateralization ratio of VVA^+^ cells in Shank3^-/-^ and Cntnap2^-/-^ mice compared to WT animals (Figs. 4A and 4B; 0.89± 0.03 in Shank3^-/-^, ANOVA, p = 0.1, n =6; in Cntnap2^-/-^: 0.88 ± 0.03, ANOVA, p = 0.08, n =5 vs. WT: 0.97± 0.02, n=7), suggesting no change in the number of BCs in the dominant somatosensory cortex. Interestingly, the lateralization ratio for the subpopulation of cells that were labeled for both VVA and PV were significantly smaller in ASD mice compared to WT mice (Fig. 4C, 0.63± 0.02 in Shank3^-/-^, ANOVA, p <0.0001, n =6; in Cntnap2^-/-^: 0.71 ± 0.06, ANOVA, p=0.0007, n =5; vs. WT: 0.97± 0.02, n=7). This was mainly due to a decrease in the proportion of PV^+^ BCs in the dominant somatosensory cortex (Fig. 4D, 57.1± 2.9% in Shank3^-/-^, ANOVA, p =0.0001, n =6; in Cntnap2^-/-^:61.02 ± 4.8%, ANOVA, p=0.0005, n =5 vs. WT: 85.2±6.4%, n=7). Together, these results suggest a slight reduction in number of BCs along with a more robust decrease in the expression of PV in BCs in the dominant hemisphere of Shank3^-/-^ and Cntnap2^-/-^ mice.

### Behavioral correlate for the decrease in PV expression in the dominant somatosensory cortex

Hypersensitivity to tactile stimulation is one of the most commonly reported symptoms in ASD (Puts NA et al. 2014). Since previous studies have shown paw dominance in mice (Waters NS and VH Denenberg 1994), we explored whether the lateralized decrease in the number of PV^+^ cells and PV expression in the dominant somatosensory cortex was correlated with tactile hypersensitivity in the dominant paw in ASD mouse models.

We thus compared sensitivity of dominant and non-dominant paws to thermal (Hargreaves test) and mechanical (von Frey) stimuli (Figs. 5 and 6). Paw withdrawal latencies (PWL) in response to thermal stimuli applied to the plantar surface of their left and right hindpaws was measured, using a Hargreaves apparatus. We measured PWLs with three different levels of heat intensities (low, medium and high intensities set at 15, 20 and 40% of maximum possible heat intensity of the Hargreaves apparatus, see Materials and Methods). A mean of four trials was calculated for each heat intensity tested. As shown in Fig.5, no differences in PWLs were found comparing dominant vs. non-dominant hindpaws of mice from all three genotypes (Fig. 5B: 15% thermal heat intensity: 2-way ANOVA, Interaction: F (2, 15)= 1.777, P=0.2030; Fig. 5C: 20% thermal heat intensity: 2-way ANOVA: Interaction: F (2, 15) = 2.368, P=0.1277; Fig. 5D: 40% thermal heat intensity: 2-way ANOVA: Interaction: F (2, 14) = 0.5479, P=0.5901). Thus, no lateralization was observed in response to heat between dominant and non-dominant paws in any of the groups. This was confirmed by calculating ratios of dominant PWL/non-dominant PWL for each paw (Fig. 5E: 2-way ANOVA: Interaction: F (4, 26) = 1.760, P=0.1672).

Next, we used von Frey filaments to apply increasing mechanical forces from weak to medium to high, using filaments of 2.83 g, 3.22 g and 4.08 g, respectively. The response to this stimulus was the percentage of withdrawals over the 2 s presentation of stimulus and was averaged for each hind paw over 10 trials for each filament. No difference was observed between the two sides with any of the three tested filaments in WT animals (Figs. 6B, 6C. and 6D.). Remarkably, we found that in both Shank3^-/-^ and Cntnap2^-/-^ mice, dominant paws were withdrawn significantly more frequently in response to weak stimulation (Fig 6B.: 2.83g: 2-way ANOVA: Interaction: F (2, 15) = 19.11, P<0.0001, Paw: F (1, 15 = 111.4), P<0.0001, Genotype: F (2, 15) = 4.44, P=0.0304) and medium (Fig 6C.: 3.22g: 2-way ANOVA: Interaction: F (2, 15) = 6.12, P<0.0114, Paw: F (1, 15 = 25.67), P=0.0001, Genotype: F (2, 15) = 2.51, P=0.1146) forces. However, with stronger mechanical stimulation, no difference was observed in responses between dominant and non-dominant paws among the three groups (Fig. 6D: 4.08g: 2-way ANOVA: Interaction: F (2, 30) = 2.735, P=0.0811). Similarly, the ratio of the responses for dominant / non-dominant paws was lateralized in these ASD mouse models (Fig. 6E: 2-way ANOVA: Interaction: F (4, 30) = 7.816, P=0.0002, Paw: F (2. 30 = 39.13), P<0.0001, Genotype: F (2, 15) = 15.70, P=0.0002). Finally, we found a strong correlation between the increase in response to low mechanical force and lateralization ratio of PV^+^ / VVA^+^ cells in each animal (R^2^=0.71, Fig. 6F). Together, these findings suggest that PV^+^ cells in the somatosensory cortex may be involved in over-reactivity to subthreshold mechanical tactile stimuli in Shank3^-/-^ and Cntnap2^-/-^ mice.

## Discussion

In this study, we provided evidence for a lateralized decrease in the number of BC interneurons that express parvalbumin (Fig. 1B and C) in the somatosensory cortex of adult Cntnap2^-/-^ and Shank3^-/-^ mice with the reduction being strongest contralateral to the dominant forepaw. This extends and is consistent with previous work showing asymmetric reductions in interneuron in other mouse models of ASD. More specifically, we trained our mice on a reaching task to identify each animal’s “dominant forepaw” and observed that the changes in interneuron numbers were more prominent in the “dominant hemisphere.” Paw sensitivity quantified using mechanical (von Frey test) stimulation showed a higher sensitivity in dominant paws compared to the other side, as well as compared to WT animals. This mechanical hypersensitivity in the dominant paw (whether right or left) strongly correlated with the ratio of the number of PV^+^ interneurons and reduced PV expression in the corresponding cortex (Fig. 6F). Such behavioral correlation provides a better understanding of the functional and clinically relevant aspects of the observed changes at the neuronal level.

Behavioral hypersensitivity of the dominant paw reported in the present study in Shank3^-/-^ and Cntnap2^-/-^ mice could simply be due to the observed lateralized decrease in PV^+^ neurons in the dominant somatosensory cortex. A recent *in vivo* population calcium imaging in the somatosensory cortex has shown increases in both spontaneous and stimulus-evoked firing in pyramidal neurons but reduced activity in interneurons resulting in sensory hyper-reactivity in Shank3^-/-^ mice(Chen Q et al. 2020). However, this study did not compare the dominant and non-dominant paw sensitivity or sensory-evoked firing properties of the two hemispheres. Recent studies have shown lateralized microstructural abnormalities in children with ASD (Peterson D *et al*. 2015) and a reduction in peak frequencies of gamma oscillations, in the dominant hemisphere during a motor task performed by the dominant index finger (An KM et al. 2018). Cortical gamma oscillations are thought to depend on PV^+^ interneurons (Cardin JA et al. 2009) and may be correlated with reduced detection thresholds. Notably, An KM et al. (2018), found that the severity of ASD symptoms was well correlated with the observed decrease in gamma oscillations. It has also been shown that a decrease in peak frequencies of gamma oscillations can be due to a decrease in PV cell numbers similar to that observed in the present study (Stroganova TA et al. 2015; Chen G et al. 2017; Geramita MA et al. 2020). Meanwhile, other factors such as dysfunction of mechanoreceptor neurons, and their connections within the spinal cord may contribute to observed behavioral hypersensitivity in our study (Orefice LL et al. 2016). Indeed a recent study has shown that mice with peripheral sensory neuron-specific deletion of Shank3 exhibit hypersensitivity in response to an air puff stimulus delivered to back hairy skin and a resultant compensatory reduction in PV^+^ neurons in S1 (Orefice LL *et al*. 2019). Here, we used von Frey test to quantify the sensitivity of glabrous skin to light touch. Although this test does not allow differentiation between central and possible peripheral dysfunction, there is no reason to think that our asymmetric behavioral responses were due to an asymmetry in the periphery. Furthermore, the above previous study has not reported an asymmetric response. Finally, our responses correlated very well with the observed anatomical asymmetry in the cortex.

About 90% of ASD individuals show deficits in processing sensory information and express abnormal responses to stimuli (Marco EJ et al. 2012). Tactile sensation is the most common sensory modality affected in ASD and the response abnormality to light touch is highly correlated with ASD severity (Tomchek SD and W Dunn 2007; Wiggins LD *et al*. 2009; Mammen MA *et al*. 2015; Orefice LL *et al*. 2019). A range of tactile abnormalities, including reduced tactile detection threshold and allodynia have been reported in children with ASD (Brambilla P et al. 2003; Abu-Dahab SM et al. 2013; Kern JK et al. 2013; Mikkelsen M et al. 2018). Here, we show an increase in detection of low intensity tactile stimuli (i.e., using 2.83g and 3.22g Von Frey filaments, which are normally considered innocuous), restricted to the dominant paw in two ASD mouse strains. This could be interpreted as lateralized allodynia. For stronger stimuli (i.e., using a stiffer Von Frey filament of 4.08g) animals showed similar response s on the two sides and similar to WT animals. These results are in line with a previous study that had shown lateralization and lower detection thresholds in response to “non-painful” tactile stimuli in the dominant hand of children with ASD (Riquelme I et al. 2016). However, both hands of these children showed reduction to the same degree in pressure pain threshold compared to typical developing children. Similar to our results, the threshold for painful stimuli was not different between two hands. While differences in the results of our study and this previous study could be due to differences in species, it is also possible that the age of subjects plays a role, that is, adult mice vs. younger children.

In our study, thermal sensitivity remained normal and without any lateralization in ASD mice. A previous study has also shown normal baseline sensitivity to thermal stimuli in Shank3^-/-^ mice (Han et al. 2016). However, impaired heat hyperalgesia was observed in inflammatory and neuropathic pain in response to injection of an inflammatory agent into the paw or a chronic constriction injury, respectively. Similar results were observed when Shank3 deletion was restricted to nociceptive peripheral neurons, which was attributed to changes in transient receptor potential subtype V1 (TRPV1) surface expression through its interaction with SHANK3. Again, this study emphasizes that not all effects are central and peripheral expression of proteins also plays an important role in generation of symptoms of ASD. However, it should be noted that the above study focused on the lower pathways (i.e., sensory terminals and spinal cord) and in general, compared to peripheral pathways, much less is known about higher processing of thermal information. One study has shown that heat sensitivity is correlated with cortical thickness (Erpelding N et al. 2012). Interestingly, although total cortical surface is different between ASD and normally developed subjects throughout life (Mensen VT et al. 2017), the cortical thickness is larger in ASD children but not in ASD adults (Khundrakpam BS et al. 2017; Mensen VT *et al*. 2017). As a result, it is possible that younger ASD mice show an abnormal response to heat stimuli.

Parvalbumin is expressed by two anatomically separate subtypes of fast-spiking cortical interneurons: chandelier cells and basket cells (Somogyi P 1977; DeFelipe J and MC Gonzalez-Albo 1998; Ariza J *et al*. 2018), which provide ∼40% of the GABAergic input to the cortex (Rudy B *et al*. 2011). Several previous studies have suggested more than 20% decrease in the number of PV^+^ interneurons in ASD individuals (Ariza J *et al*. 2018; Hashemi E *et al*. 2018) and mouse models (Sadakata T *et al*. 2007; Selby L et al. 2007; Gogolla N *et al*. 2009; Tripathi PP *et al*. 2009; Zikopoulos B and H Barbas 2013), and others have found a reduction in PV expression in cortical BCs without an overall reduction in the number of these cells (Filice F *et al*. 2016; Lauber E *et al*. 2018). In the present study, we found a decrease in both the number PV^+^ BCs and the expression of PV in BCs without any change in overall number of BCs. Simply considering percent changes in these values suggests a more prominent effect on PV expression than cell number in both Cntnap2^-/-^ and Shank3^-/-^ mice. More significantly, we found that this reduction in PV expression was not random and was more robust in the dominant hemisphere. However, since we did not study chandelier cells, we are not able to exclude any concomitant changes in PV^-^ chandelier cells in the somatosensory cortex of these mice. A recent study reported a reduction in chandelier cells in the medial prefrontal cortex of several individuals with ASD (Ariza J *et al*. 2018). This conclusion was based on histology results that showed a decrease in the number of PV^+^ and VVA^-^ cells, which represent chandelier cells. However, it is also not clear whether these cells have lost their PV expression (similar to that shown here) or that the cells did undergo atrophy and death. Also, it should be noted that only 15% of chandelier cells in the medial prefrontal cortex are PV^+^ (Taniguchi H et al. 2013).

In addition to PV^+^ neurons, two other groups of interneurons exist that can be identified by the presence of SST or 5HT3a receptors. Developmentally, cortical interneurons arise from either medial (MGE) or caudal (CGE) ganglionic eminences. Additionally, lateral ganglionic eminences (LGE) is the primary source of olfactory bulb interneurons (Batista-Brito R and G Fishell 2009). CGE interneurons express 5HT3a and MGE interneurons express PV or SST (Kelsom C and W Lu 2013). Our results regarding MGE derived total cell counts in ASD confirm findings of previous studies in ASD models that have shown a decrease in PV^+^ cells (Gogolla N *et al*. 2009; Penagarikano O *et al*. 2011; Ariza J *et al*. 2018; Hashemi E *et al*. 2018) and no change in SST^+^ cells (Speed HE et al. 2015; Vogt D *et al*. 2018). Here, we also investigated changes in CGE derived interneurons, which express 5-HT3a, since disruption of the 5-HT system during early development has been linked to impaired autistic-like social behavior (Engel M et al. 2013). Surprisingly, we did not find any changes in the number of cells with 5-HT3a or its expression by interneurons in Shank3^-/-^ and Cntnap2^-/-^ mice. Our results show that the above mutations only affect PV expressing MGE derived neurons. The reduced migration probably underlies the lower cell counts and the lower expression of PV can be due to inadequate activity in PV cells. The expression of PV in BCs around P10 – P14 partly depends on neural activity and develops at different postnatal times and rates in the left and right hemispheres (Thatcher RW et al. 1987). Thus, any disruption in the activity in one hemisphere during this critical period can result in a unilateral decrease in PV expression, as observed in our ASD mice. PV is a slow Ca^2+^ binding protein that affects the amplitude and time course of presynaptic intracellular Ca^2+^ transients after an action potential and affects short term plasticity (Caillard O et al. 2000). Expression of parvalbumin in interneurons of the cerebellum, hippocampus and striatum, contributes to presynaptic short-term facilitation (Caillard O *et al*. 2000; Vreugdenhil M et al. 2003). Therefore, reduction in PV expression in layer 2/3 can result in short term facilitation that will affect the inhibitory inputs of these interneurons on both inhibitory and excitatory cells in the circuit. Based on the behavioral results presented here, our prediction is that the net effect is an increase in excitation. This can be explained at least in part by the loss of PV^+^ interneurons (in addition to loss of PV expression).

It is important to note that minimal alteration in the number of PV^+^ interneurons as seen in the present study can potentially have a dramatic impact on cortical microcircuits and their function (DeFelipe J 1999; Ariza J *et al*. 2018). Each chandelier cell or BC innervates up to 200 pyramidal cells. BCs affect the output of networks of pyramidal cells by innervating their soma and proximal dendrites (Halasy K et al. 1996; Czeiger D and EL White 1997; Szabo GG et al. 2014). A similar interhemispheric imbalance in the number of PV^+^ cells has also been reported in NL-3^-/-^ and VPA mouse models of ASD (Gogolla N *et al*. 2009). However, this imbalance was not linked to the dominant hemisphere or any behavioral changes. Together, our findings along with the results of previous studies suggest an alteration in inhibitory circuits as a unifying neuronal mechanism between different types of ASD.

Finally, a systematic correlation has been shown by previous studies between handedness and hemispheric language dominance (Knecht S et al. 2000). Although we focused on the lateralization in PV^+^ cells in the somatosensory cortex, we observed similar changes in other primary sensory areas, particularly in the auditory cortex (data were not shown). Processing auditory information and perceiving the meaning of sounds is a critical component of verbal communication, and shows deficits in ASD (Moore BC et al. 2008; Marco EJ et al. 2011; Tietze FA et al. 2019). For example, fMRI in individuals with Asperger’s syndrome shows weaker activation in the dominant auditory hemisphere compared to typically developing control. It has been suggested that such alterations in auditory processing in the dominant hemisphere, underlies the difficulties in speech perception in Asperger’s syndrome (Tietze FA *et al*. 2019). Future detailed study may reveal the possible involvement of lateralization of PV cells in the dominant auditory cortex of ASD individuals and their language development.

---

**Supplementary Figure 1.**
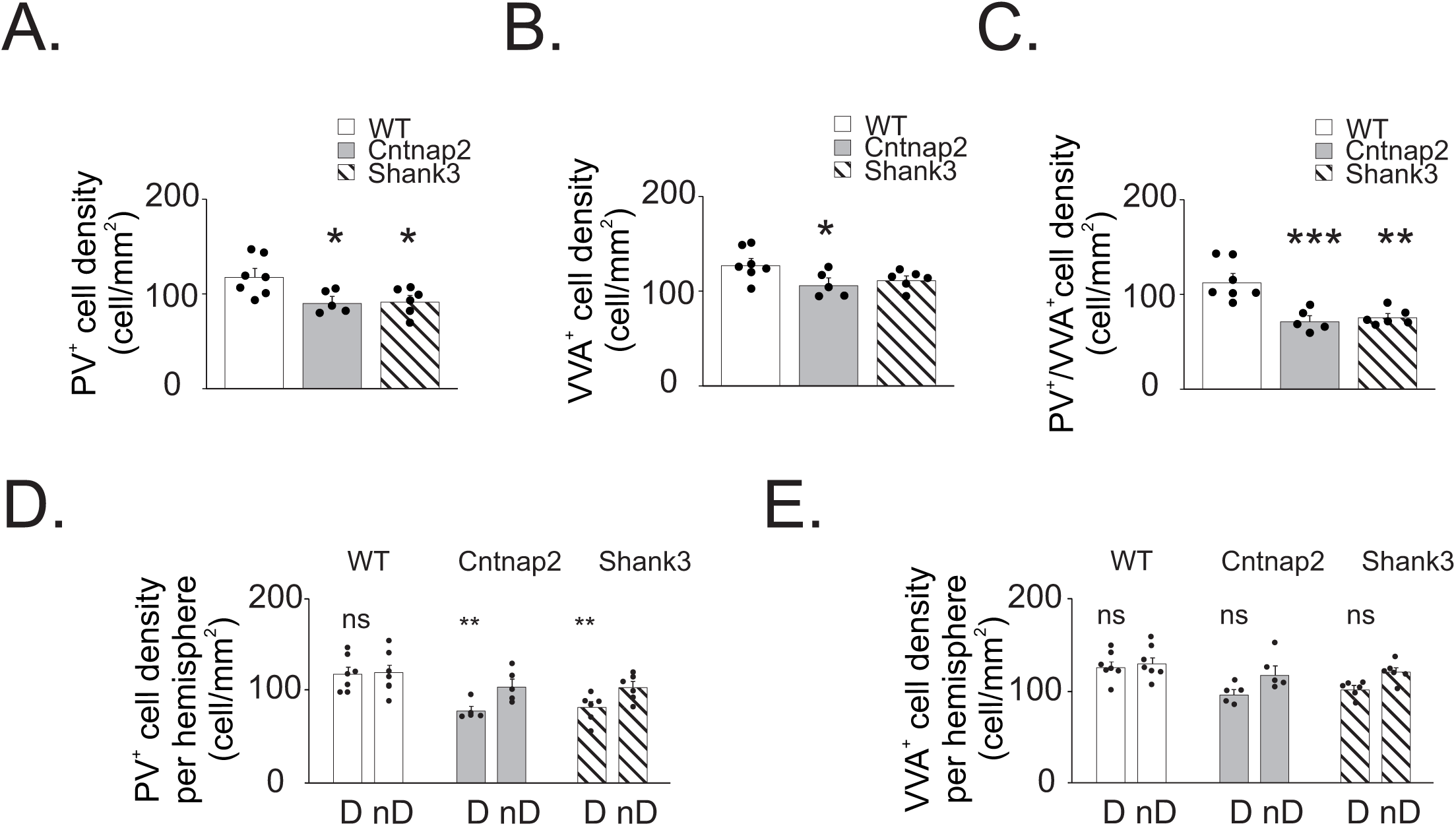

**Supplementary Figure 2.**
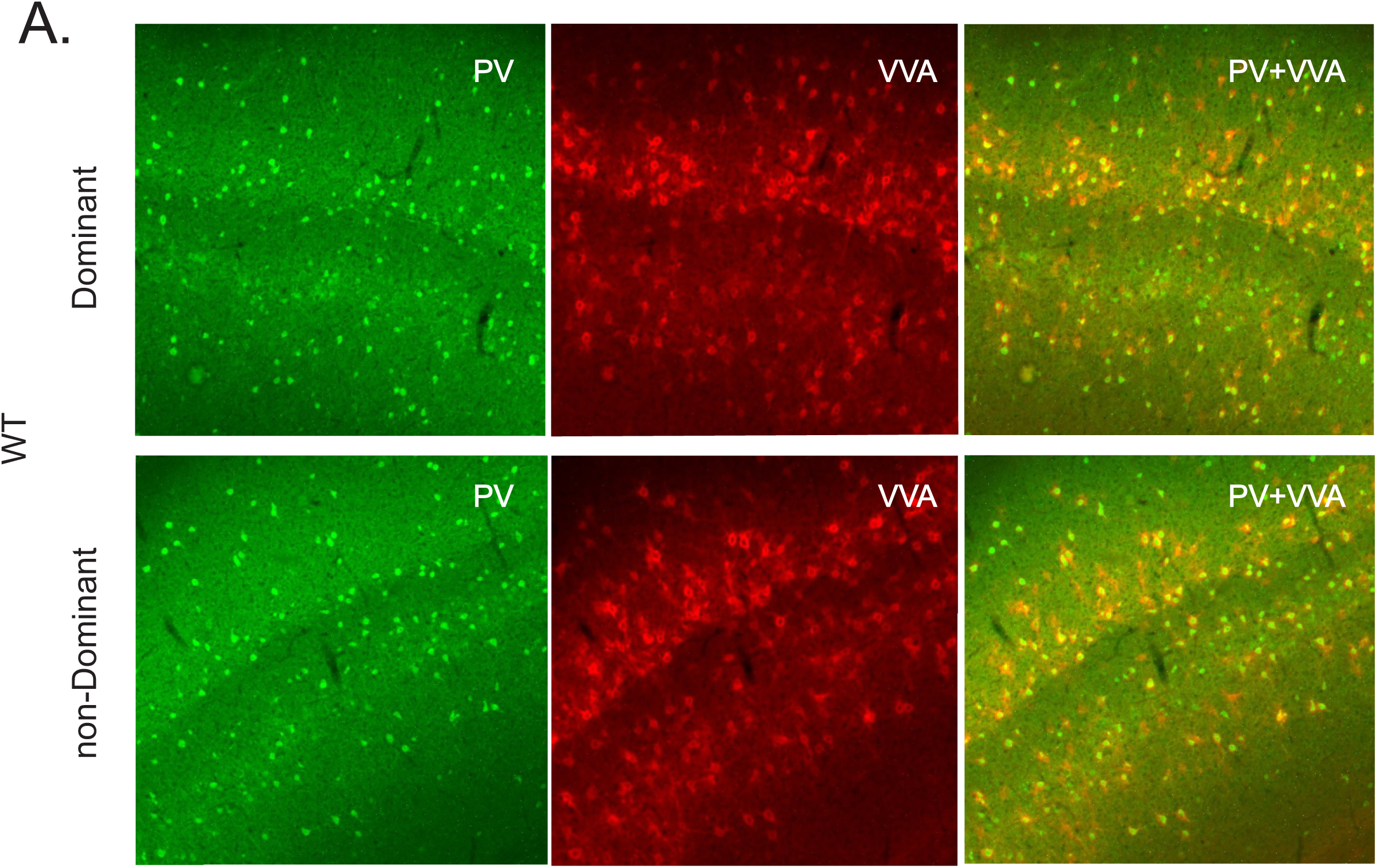

